# The new generation hDHODH inhibitor MEDS433 hinders the *in vitro* replication of SARS-CoV-2

**DOI:** 10.1101/2020.12.06.412759

**Authors:** Arianna Calistri, Anna Luganini, Valeria Conciatori, Claudia Del Vecchio, Stefano Sainas, Donatella Boschi, Marco Lucio Lolli, Giorgio Gribaudo, Cristina Parolin

## Abstract

Identification and development of effective drugs active against SARS-CoV-2 are urgently needed. Here, we report on the anti-SARS-CoV-2 activity of MEDS433, a novel inhibitor of human dihydroorotate dehydrogenase (hDHODH), a key cellular enzyme of the *de novo* pyrimidines biosynthesis. MEDS433 inhibits *in vitro* virus replication in the low nanomolar range, and through a mechanism that stems from its ability to block hDHODH activity. MEDS433 thus represents an attractive candidate to develop novel anti-SARS-CoV-2 agents.

## Main text

The emergence of the novel Severe Acute Respiratory Syndrome Coronavirus 2 (SARS-CoV-2), and the rapid worldwide spreading of coronavirus disease 19 (COVID-19) have produced a threat to global public health that calls for urgent deployment of effective antiviral drugs (1–4). Among the therapeutic options that have been potentially considered, small-molecules targeting host factors exploited by SARS-CoV-2 to replicate may represent an alternative to direct acting agents prone to select drug resistant strains (5). One of the cellular pathways that is attracting more attention for the advancement of host-targeting antivirals (HTA), is the *de novo* pyrimidines biosynthesis, essential for virus replication in infected cells (6). In this pathway, the human dihydroorotate dehydrogenase (hDHODH) catalyzes the rate-limiting step of dehydrogenation of dihydroorotate to orotate, thus providing uridine and cytidine to fulfill nucleotides request (7–9). Given its critical role, hDHODH is considered an emerging target of choice for the development of HTA against SARS-CoV-2 (10). In this regard, two potent hDHODH inhibitors just entered in Phase II clinical trials for COVID-19: brequinar (11) (NCT04425252), and PTC299 (12) (NCT04439071). However, these drug candidates suffer of toxicity issues that slowed down their earlier clinical pathway on other therapeutic applications (13, 14). Thus, new safest hDHODH inhibitors are urgently needed.

To investigate the feasibility of targeting hDHODH to develop HTA against SARS-CoV-2, in this study we have characterized the *in vitro* antiviral activity of a new generation hDHODH inhibitor produced by the rational modulation to brequinar (11), and characterized by the presence of a 2-hydroxypyrazolo[1,5-a]pyridine moiety. This compound, MEDS433 (Fig. 1A), is comparable to brequinar in inhibiting *h*DHODH activity, while owing a better *drug-like* profile (15, 16).

**Figure 1.**
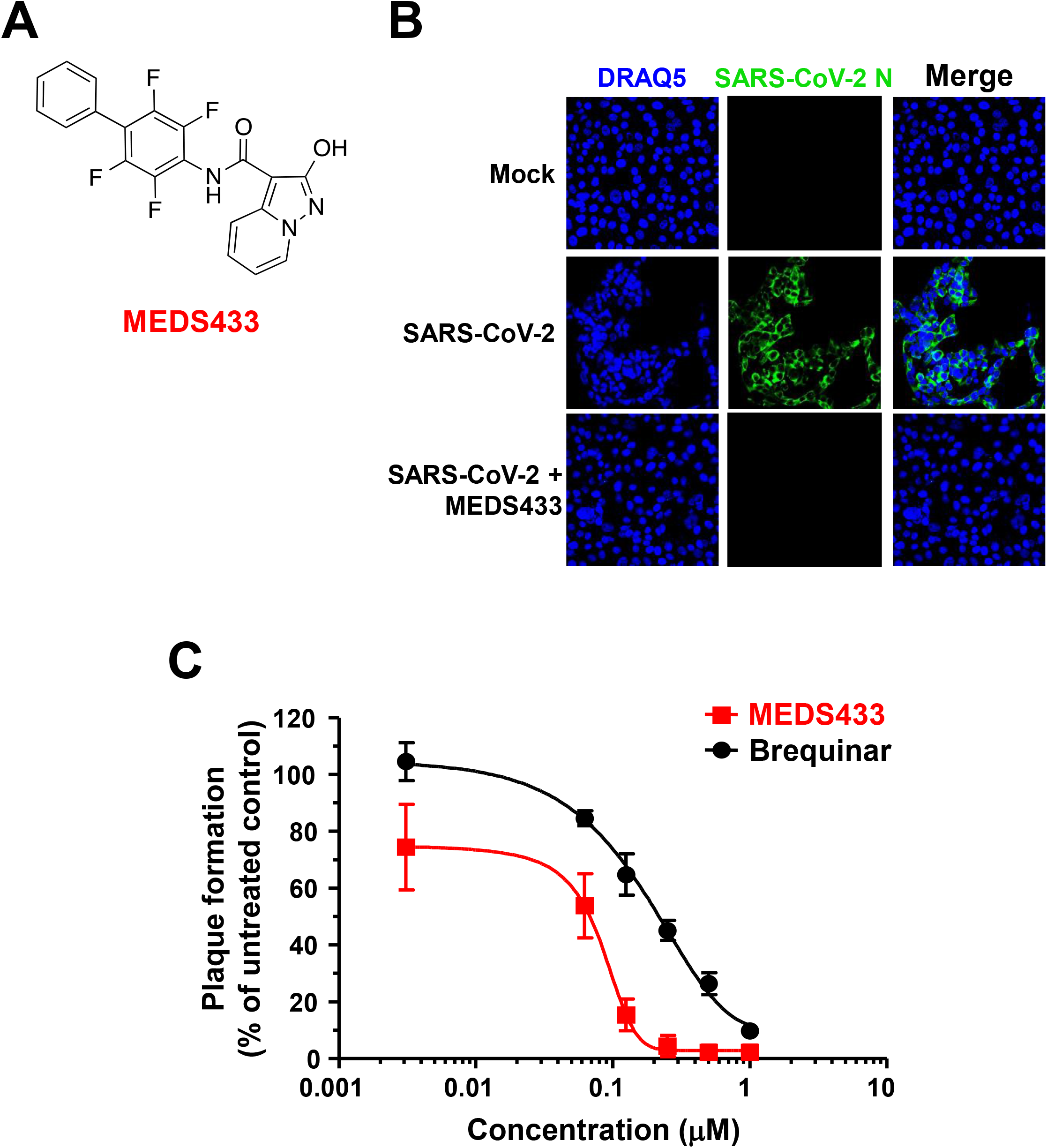
Antiviral activity of MEDS433 on SARS-CoV-2 replication. **(A)** Structure of MEDS433. **(B)** Immunofluorescence analysis of SARS-CoV-2-infected cells. Vero E6 cells were treated with vehicle (DMSO) or with 0.5 μM MEDS433 1 h prior to infection with SARS-CoV-2 at an MOI of 0.1. At 24 h p.i., cells were fixed, permeabilized, and immunostained with an anti-SARS-CoV-2 nucleocapsid protein (N) mAb, followed by Alexa 488-conjugated secondary antibody. Nuclei were stained with DRAQ5. Confocal laser microscopy images acquired in the green (SARS-CoV-2 N) and the blue (DRAQ5) channels are shown, as well as overlaid images (merge). **(C)** Dose dependent inhibition of SARS-CoV-2 replication by MEDS433. Vero E6 cell monolayers were infected with SARS-CoV-2 (50 PFU/well), and, where indicated, the cells were treated with vehicle (DMSO) or increasing concentrations of MEDS433 or brequinar 1 h before, during virus adsorption, and throughout the experiment. At 48 h p.i., infectious SARS-CoV-2 in cell supernatants was titrated by plaque assay on Vero E6 cells. MEDS433 and brequinar concentrations producing 50 and 90% reductions in plaque formation (EC_50_ and EC_90_, respectively) were determined as compared to control treatment (DMSO). The data shown represent means ± SD (error bars) of three independent experiments performed in triplicate.

The effect of MEDS433 on SARS-CoV-2 replication was evaluated in Vero E6 cells infected with a clinical isolate of SARS-CoV-2 (2019-nCoV/Italy-INMI1) at a multiplicity of infection (MOI) of 0.1, and then treated with 0.5 μM MEDS433. At 24 hours post-infection (h p.i.), cells were fixed, permeabilized and stained with an anti-SARS-CoV-2 nucleocapsid N protein mAb, and with DRAQ5 which stains cell nuclei, to count cell numbers. As shown in Fig. 1B, confocal microscopy revealed that while about 85% of infected control cells expressed the N protein, MEDS433 treatment completely abolished its accumulation, thus indicating that N protein expression could be prevented by targeting the *de novo* pyrimidine biosynthesis.

Next, a virus yield reduction assays (VRA) was performed in SARS-CoV-2-infected Vero E6 cells treated with increasing concentrations of MEDS433. At 48 h p.i., cell supernatants were harvested and titrated by plaque assay. A concentration-dependent inhibition of SARS-CoV-2 replication was thus observed (Fig.1C), with EC_50_ and EC_90_ values of 0.063 and 0.136 μM, respectively. Interestingly, MEDS433 was more effective than brequinar (EC_50_ 0.20, EC_90_ 1μM) (Fig. 1C). In addition, the Cytotoxic Concentration (CC_50_) of MEDS433 as measured in uninfected Vero E6 cells by the MTT method (17), was more than 500 μM with a favorable Selective Index (SI) greater than 7,900, thus indicating that its antiviral activity was not due to a reduced cell viability.

To get more insights into MEDS433 mechanism of action, time-of-addition experiments were carried out. Briefly, Vero E6 cells were exposed to MEDS433 (0.5 μM) from −2 to - 1h prior to SARS-CoV-2 adsorption (MOI of 0.1) (pre-treatment); during infection (adsorption stage, from −1 to 0 h) (co-treatment); or after viral adsorption (from 0 to 48 h p.i.) (post-treatment). Infectious SARS-CoV-2 particles were then quantified in cell supernatants harvested at 48 h p.i. by plaque assay. As depicted in Fig. 2A, MEDS433 did not affect the initial attachment and entry phases of the SARS-CoV-2 life cycle, while it produced a significant reduction of infectious virus production when added at a post-entry stage, in agreement with its ability to block N protein accumulation (Fig. 1B). Immunoblot analysis of total protein extracts prepared from the corresponding SARS-CoV-2-infected- and MEDS433-treated cells and fractionated through a 10% SDS-PAGE, confirmed a reduction of N protein content only in the post-treatment sample (Fig. 2B), thus indicating that MEDS433 interferes with a post-entry biosynthetic step in SARS-CoV-2 replication.

**Figure 2.**
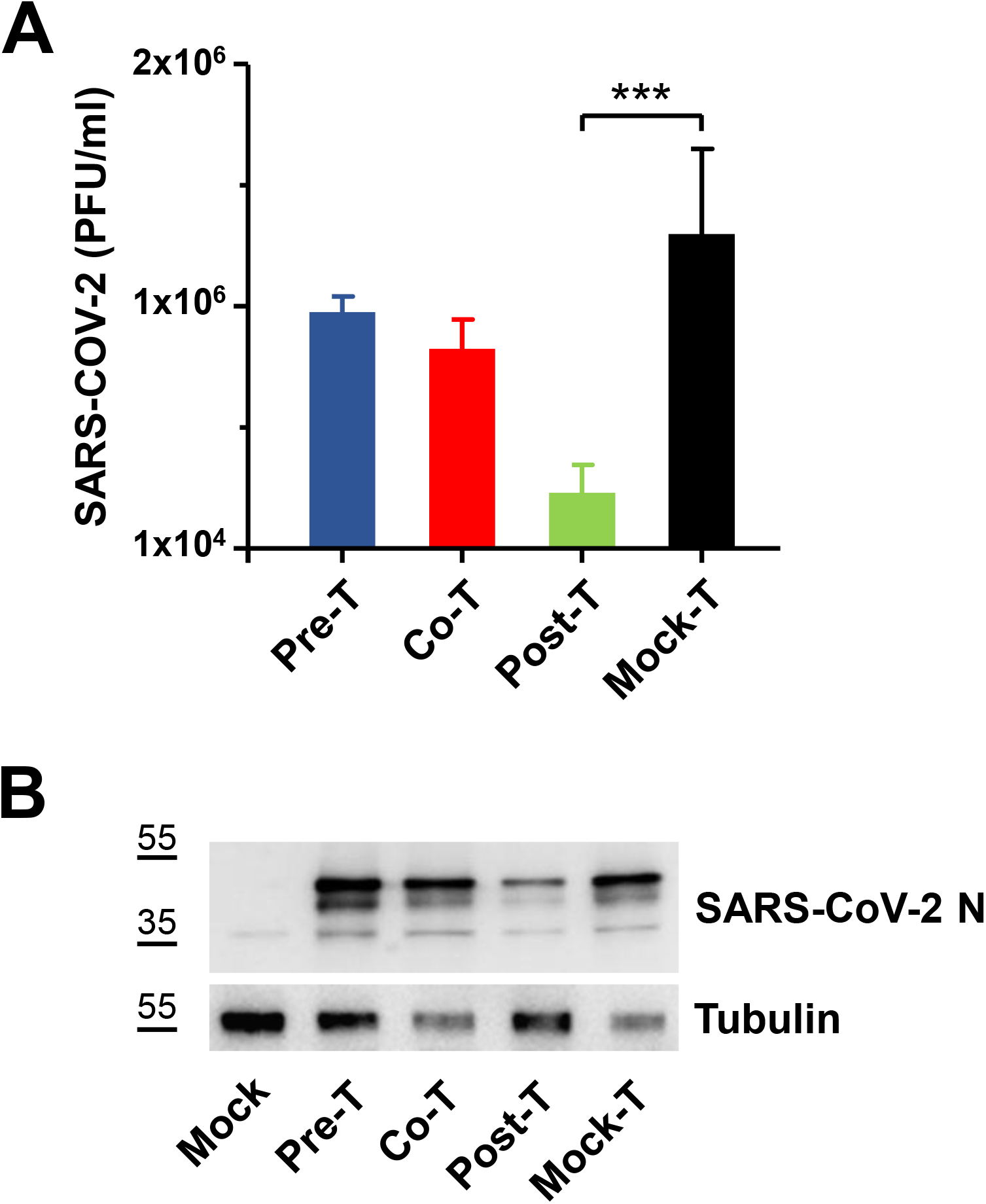
MEDS433 targets a post-entry stage in the SARS-CoV-2 replicative cycle. **(A)** Time-of-addition experiment. Vero E6 cells were incubated with vehicle (DMSO) or with 0.5 μM MEDS433 from 2 to - 1h prior to SARS-CoV-2 infection (MOI of 0.1) (pre-treatment, Pre-T); during infection (from −1 to 0 h) (co-treatment, Co-T); or after viral infection (from 0 to 48 h p.i.) (post-treatment, Post-T). Thereafter, production of infectious SARS-CoV-2 was measured by titrating cell supernatants by plaque assay on Vero E6 cells. The data shown represent means ± SD (error bars) of three independent experiments performed in triplicate. Statistical significance was calculated by a one-way ANOVA followed by Dunnett’s multiple comparison test. *** (p < 0.0001) compared to the calibrator sample (MEDS433 alone). **(B)** Immunoblot analysis of SARS-CoV-2 nucleocapsid protein. Total protein extracts prepared from SARS-CoV-2-infected Vero E6 cells monolayers treated as described in (A), were fractionated through a 10% SDS-PAGE, and immunoblotted with an anti-N protein mAb. Tubulin was immunodetected as protein loading control.

These results suggested a mechanism of the anti-SARS-CoV-2 activity of MEDS433 consistent with the hypothesis of an interference with the *de novo* pyrimidine biosynthesis. To verify this hypothesis, we investigated by plaque reduction assays (PRA) in Vero E6 cells whether the antiviral activity of MEDS433 could be overcome by supplementing cell medium with increasing concentrations of exogenous uridine to bypass the requirement of *de novo* pyrimidine biosynthesis. As shown in Fig. 3 (upper panel), the anti-SARS-CoV-2 activity of 0.3 μM MEDS443 was significantly reversed by a 100-fold excess of uridine relative to MEDS433 concentration, and completely overturned by greater uridine concentrations, thus confirming that the *de novo* pyrimidine pathway was inhibited by MEDS433 in SARS-CoV-2-infected cells. Then, to conclusively prove that hDHODH inhibition was responsible of MEDS433 antiviral effect, increasing concentrations of the hDHODH substrate dihydroorotic acid or its product, orotic acid were added to cell medium. In SARS-CoV-2-infected Vero E6 cells treated with MEDS433 (0.3 μM), the addition of orotic acid reversed in a dose dependent manner the antiviral effect of MEDS433 (Fig. 3, lower panel), with complete reversion observed at the highest concentration (1000 x the MEDS433 concentration) evaluated. In contrast, the supplement of dihydroorotic acid, even at 1 mM (3,333 times more than MEDS433), did not affect MEDS433 antiviral activity (Fig. 3, lower panel), thus indicating that MEDS433 inhibited a step in the *de novo* pyrimidine biosynthesis pathway downstream from dihydroorotic acid. Altogether, these results clearly confirmed that MEDS433 specifically targets hDHODH activity in SARS-CoV-2-infected cells, and that this inhibition is responsible of its overall antiviral activity.

**Figure 3.**
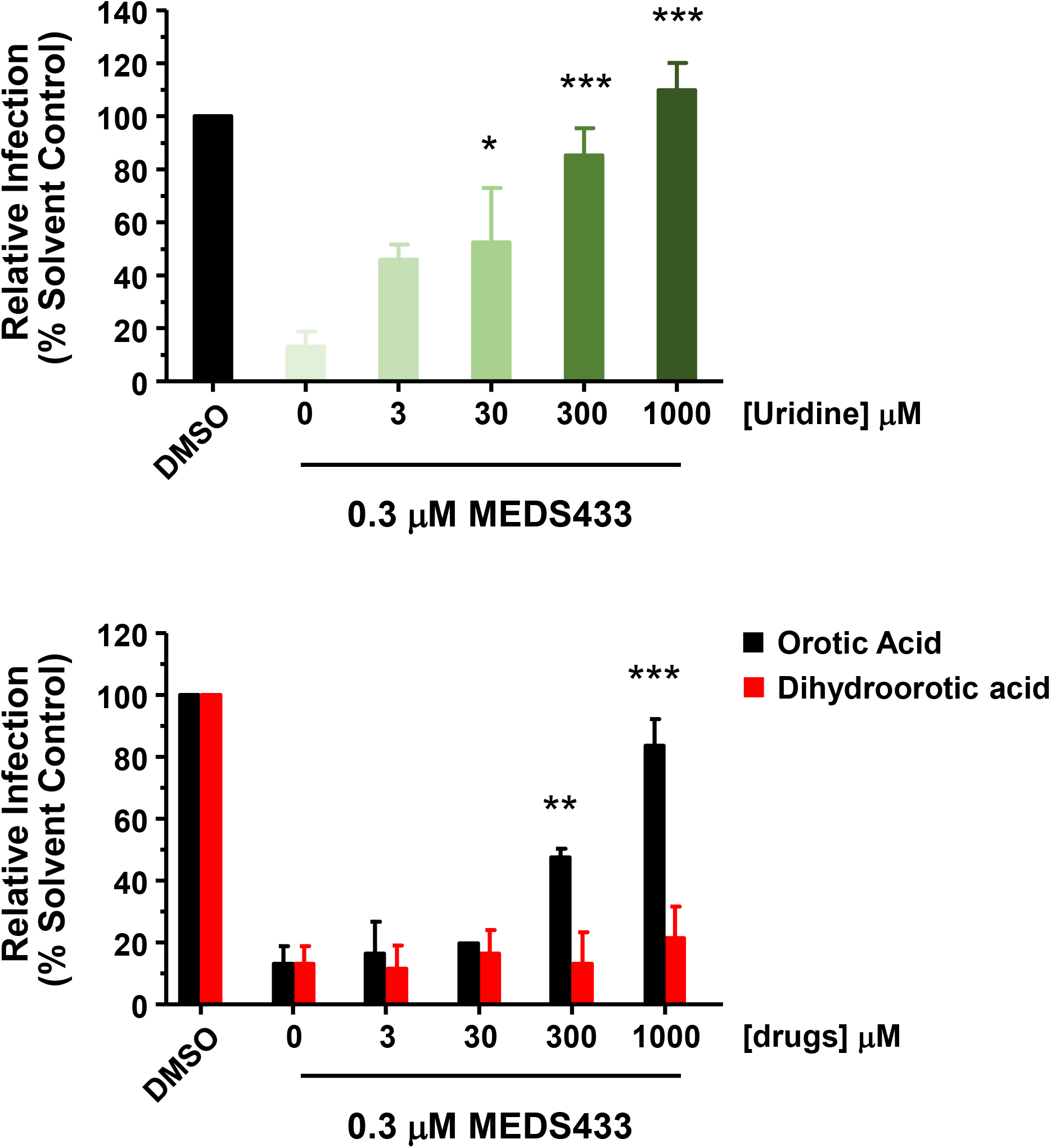
Uridine or orotic acid supplementation counteracts the anti-SARS-CoV2 activity of MEDS433. Vero E6 cell were treated with solvent (DMSO) or 0.3 μM of MEDS433 in the absence or presence of increasing concentrations of uridine (upper panel), orotic acid or dihydroorotic acid (lower panel) before and during infection with SARS-CoV-2 (30 PFU/well). Following virus adsorption, compounds were added to cell monolayers and viral plaques were then stained and were microscopically counted at 48 h p.i.. Plaque counts for each drug concentration were expressed as a percent of the mean count of the control cultures treated with DMSO. The data shown represent means ± SD of three independent experiments performed in triplicate. Statistical significance was calculated by a one-way ANOVA followed by Dunnett’s multiple comparison test. ** (p < 0.0001), ** (p < 0.001) and * (p < 0.05) compared to the calibrator sample (MEDS433 alone).

In conclusion, our study while confirming along with others recent reports (12,18) hDHODH as a noteworthy target to inhibit SARS-CoV-2 infection (10,19), highlights MEDS433 as an attractive candidate to develop HTA for COVID-19. MEDS433 in fact performs better than brequinar for both antiviral potency and SI (this study and 16), and its safety profile is superior to that of PTC299 (12). Therefore, the potent *in vitro* anti-SARS-CoV-2 activity of MEDS433 and its valuable *drug-like* profile, support further studies to validate its therapeutic efficacy in preclinical animal models of COVID-19.

## Acknowledgments

This work was supported by Italian Ministry for Universities and Scientific Research (Research Programs of Significant National Interest, PRIN 2017–2020, Grant No. 2017HWPZZZ_002) to AL.; Ministero degli Affari Esteri e della Cooperazione Internazionale (Grant number PGR01071 Italia/Svezia (MIUR/MAECI)) to D.B. and M.L.L.; Associazione Italiana per la Ricerca sul Cancro (AIRC) Individual Grant 2019 (AIRC IG 2019 DIORAMA 23344) to D.B. and M.L.L.; the University of Torino (Ricerca Locale) to D.B., G.G., M.L.L. and A.L.; the University of Padua (DOR) to A.C. and C.P.; PARO_FINA20_01 to C.P.; and BIRD grant CALI_SID19_08 to A.C. This study was also supported by the European Virus Archive goes Global (EVAg) project that has received funding from the European Union’s Horizon 2020 research and innovation programme under grant agreement No 653316.

## Notes

### Competing Interest Statement

The authors have declared no competing interest.

